# Time-Space Resolved Fluorescence Spectroscopy in Live *Chlamydomonas* Cells under Light-Harvesting Regulation

**DOI:** 10.1101/2024.10.10.617661

**Authors:** Yuki Fujita, Xianjun Zhang, Shen Ye, Yutaka Shibata

## Abstract

We report here a technical advancement that enables time-resolved fluorescence spectroscopy in spatially resolved domains of a living cell at low temperatures. The technique is based on a combination of the self-developed cryo-confocal microscope system and the streak-camera technology. An instrumental response time of ca. 24 ps was achieved. This technique was applied to reveal the light-harvesting dynamics in local domains within single *Chlamydomonas reinhardtii* cells. Organisms performing oxygenic photosynthesis, like *Chlamydomonas*, have evolved a regulation mechanism called state transitions (ST), which maintains the excitation balance between PSI and PSII. ST relies on the shuttling of light-harvesting chlorophyll protein complex II (LHCII) between the two PSs. In the present experiment, cells were induced either to state1, where LHCII is bound to PSII, or state2, where LHCII moved and is bound to PSI. After the induction of ST, cells were immediately cooled to ca. 80 K, where PSI and PSII show clearly separated fluorescence emission bands, enabling the visualization of these components separately. Based on kinetic analyses of the time-resolved fluorescence spectra in both PSI-rich and PSII-rich local domains, we concluded that (1) the intracellular inhomogeneity in the PSII/PSI fluorescence ratio comes from that in the PSII/PSI stoichiometry, not from that in the antenna sizes of the PSs, and (2) the antenna size of PSI in state2 cells may larger in intact cells than that of the isolated PSI-LHCI-LHCII super-complex reported so far.

## Introduction

Oxygenic photosynthesis is performed by two photosystems (PSs), called PSII and PSI. These two PSs drive the linear electron flow from water to nicotinamide adenine dinucleotide phosphate (NADP^+^) by using the energy of sunlight. Thus, they can be regarded as tandem solar cells. Maintaining the excitation balance between the two PSs is important for efficient linear electron flow from water to NADP^+^. Oxygenic photosynthetic organisms have developed a physiological function, called state transitions (ST), to realize the above-mentioned maintenance.(Delosme et al., 1996; Minagawa and Tokutsu, 2015) Its mechanism is based on the shuttling of the light-harvesting chlorophyll protein complex II (LHCII) between the two PSs. Here, LHCII is one of the peripheral antennas feeding the excitation energy to PSs. LHCIIs are bound to PSII to feed the excitation energy under ordinary conditions. This state is called the state1. When the environmental light prefers to excite PSII more, LHCIIs are detached from PSII and bind to PSI. This is called state2.

Many biochemical and structural-biological studies have provided evidence for the migration of LHCII upon the ST.(Takahashi et al., 2006; Tokutsu et al., 2009; Snellenburg et al., 2017; Huang et al., 2021; Pan et al., 2021) On the other hand, there have been only limited reports on the visualization of the LHCII movement by the optical microscopic observation of living cells.(Iwai et al., 2010; Kim et al., 2015; Fujita et al., 2018; Zhang et al., 2021; Fujita et al., 2022; Zhang et al., 2022; Verhoeven et al., 2023) Such approaches will be important for clarifying the dynamic aspect of the ST. They specifically allow observation of the intermediate state of the ST, where LHCIIs are not yet bound to either of the PSs but are drifting in the space between the two PSs. The characterization of the isolated free LHCII arising during the ST is an important issue to be clarified. We have made an effort in this direction by using the self-developed cryo-spectral microscope system.(Shibata et al., 2014; Fujita et al., 2018; Fujita et al., 2022) The low-temperature fluorescence spectrum of an oxygenic photosynthetic organism usually shows clearly separated bands assigned to PSII at around 680-695 nm and to PSI at around 710-730 nm. The specifically red-shifted fluorescence band of PSI at low temperatures is due to the so-called red chlorophylls (Chls), which are those with excitation energies well below that of P700, the primary electron donor of PSI. Owing to the characteristic separation of the PSI and PSII fluorescence bands,(Lamb et al., 2018) the measurement of the fluorescence spectra under the microscope at a low temperature is effective in resolving the intracellular PSI-rich and PSII-rich domains.(Fujita et al., 2018; Chiba and Shibata, 2019; Fujita et al., 2022) Furthermore, detailed analyses of the spectral features provide information on the peripheral antenna.

Although the cryo-microscopic measurements of the ST in intact cells are effective as described above, there have been several difficulties in obtaining insight into the state of LHCs during the ST based only on the intracellular local fluorescence spectra. Since the fluorescence intensity is proportional to both the concentration and the fluorescence quantum yield of the emitter, there remains ambiguity in determining the state of the LHC. We have extended the cryo-microscopic system to enable the simultaneous detection of the local fluorescence spectra and decay kinetics of 680-emitting species (fluorescence lifetime imaging: FLIM)(Fujita et al., 2022) to determine both the fluorescence spectra and quantum yield. This was a great advancement. However, the fluorescence decay kinetics could be detected only at one single wavelength with the developed FLIM system, which still limited our ability to clarify the local state of the antenna within single cells. Based on the fluorescence kinetic trace only at one wavelength, we could not determine whether a very rapid fluorescence decay originated from an efficient excitation-energy transfer (EET) process or a rapid quenching. There is another troubling problem in evaluating PSI-rich and PSII-rich local domains within single cells. Though the local PSII/PSI fluorescence intensity ratio was utilized to evaluate the PSII-rich or PSI-rich domains, in principle, the ratio can be affected by an inhomogeneity in the antenna size of the PSs. This situation is explained in Fig. 1. The cryo-FLIM did not provide any clear answer to this question.

**Figure 1.**
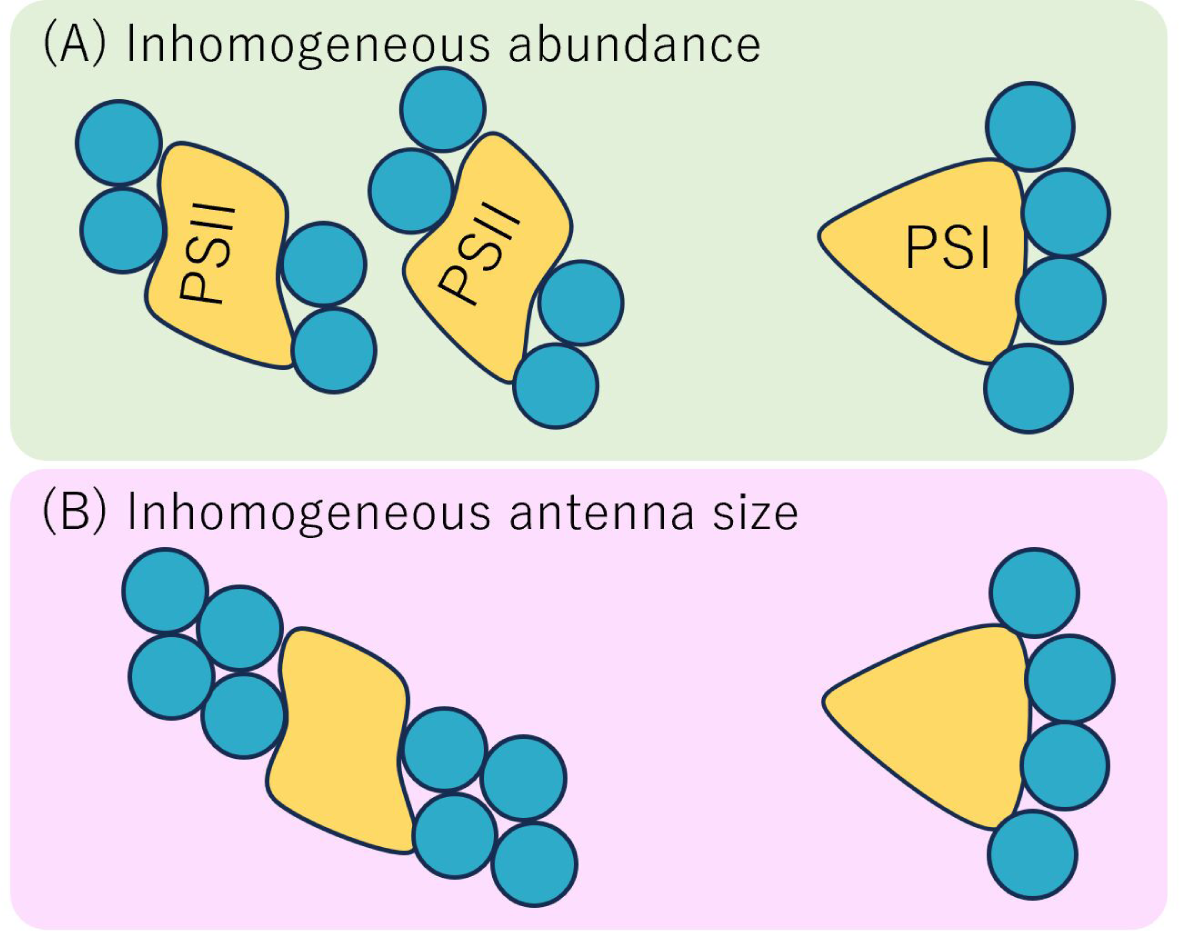
Schematic description of two possible situations resulting in high relative local PSII fluorescence. It is assumed that the spatially inhomogeneous PSII fluorescence originates from (A) inhomogeneous concentrations of PSs and (B) inhomogeneous antenna size. Blue and green circles are the peripheral antennae associated with PSII and PSI, respectively.

In the present study, we further extended the cryo-microscopic system to enable measurement of the time-resolved fluorescence spectra over the whole spectral range of the chlorophyll (Chl) fluorescence in the local PSII-rich and PSI-rich domains separately. This was achieved by connecting the cryo-microscopic system with the streak camera together with updating the control program. The developed system enabled the local time-resolved fluorescence spectra with an instrumental response time of ca. 24 ps. This advancement allowed us to conduct detailed analyses of the light-harvesting dynamics in the local domains within intact cells. Here, we demonstrated the implementation of this updated setup. We investigated the rapid fluorescence kinetics of photosynthetic components upon the operation of ST of a unicellular green alga, *Chlamydomonas reinhardtii*, that has been a model organism of the photosynthetic study. We could evaluate the EET dynamics of the PSI-rich and PSII-rich domains separately. Furthermore, we emphasize that the present technical advancement will provide a powerful method for analyzing a wide variety of research targets, including not only the natural photosynthetic system but various artificial systems as well.

## Results

Figure 2 shows a typical result of the first raster scanning detected by the polychromator-CCD camera system. Panels (A) and (B) show, respectively, the fluorescence image and the PSII/PSI ratio map of a *Chlamydomonas* cell in state1. The image in Fig. 2A shows a cup-shaped chloroplast, which is typical of *Chlamydomonas*. It wraps the nucleus corresponding to the dark area on the left-hand side of the cell. The structure of a subcellular organelle called pyrenoid can be found on the right-hand side. Pixels shown in blue and red in Fig. 2C are those assigned to the PSII-rich and PSI-rich domains with the *R*_PSII_and *R*_PSI_values beyond the thresholds, respectively. Fluorescence spectra averaged over the PSII-rich and PSI-rich domains are shown by the blue and red curves in Fig. 2D, respectively.

**Figure 2.**
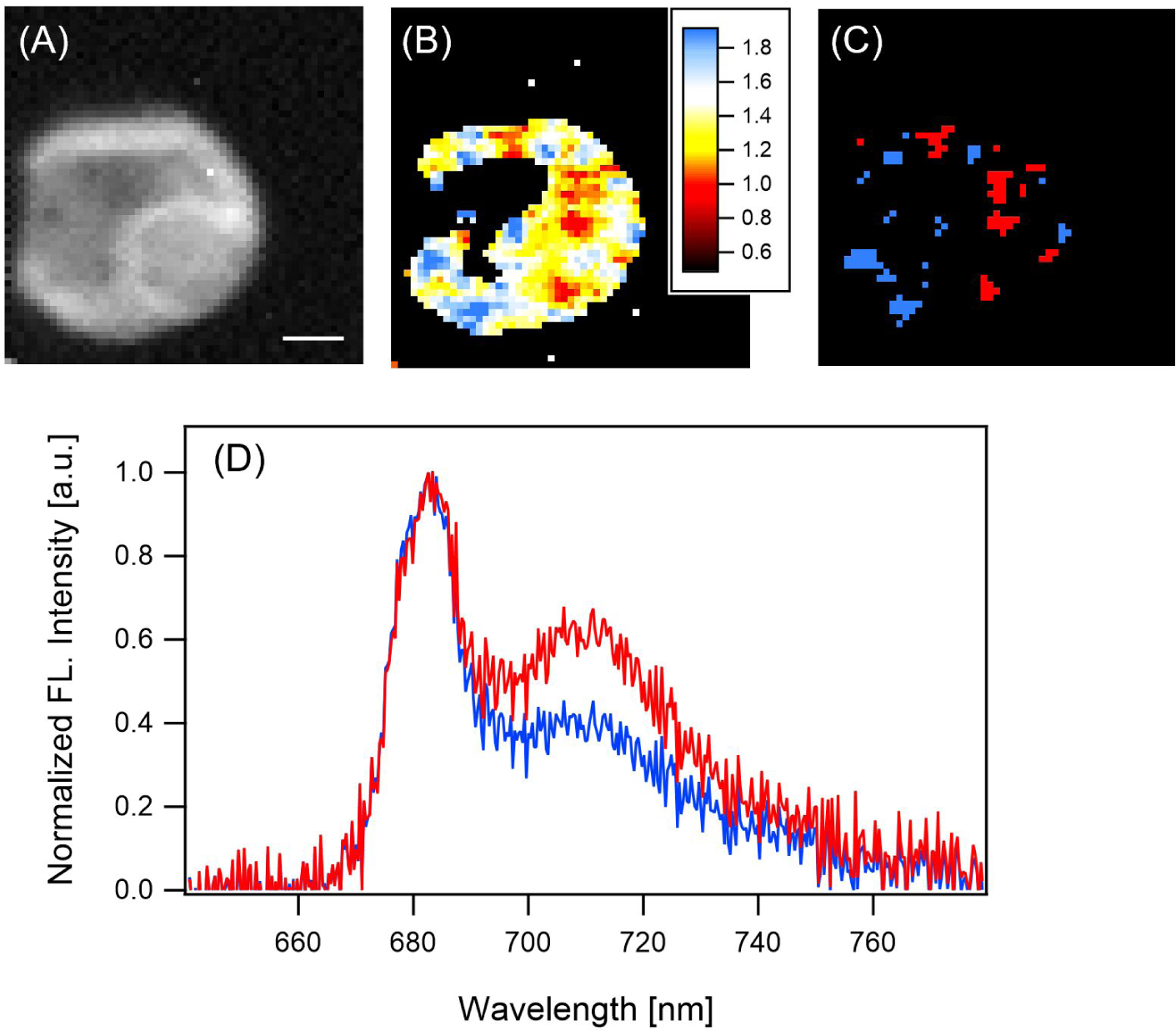
(A) A typical fluorescence image of a *Chlamydomonas* cell induced to state1 detected by the CCD camera. The scale bar indicates 2 µm. (B) PSII/PSI ratio map regenerated by dividing the fluorescence intensity integrated over the range of 676-692 nm (PSII preferential wavelength) by that over the range of 706-727 nm (PSI preferential wavelength) for every pixel of panel (A). (C) Blue and red fills highlight 50 pixels with the largest and smallest PSII/PSI values, respectively. (D) Blue and red solid lines represent normalized fluorescence spectra averaged over the regions specified by blue and red pixels in panel (C).

The second and third scans were done only over the PSII- and PSI-rich domains, respectively. The signal acquisitions of the streak-camera system were conducted during the second and third selective scans. In this case, we set the dwell time at each pixel to 5 s to increase the signal intensity. Since the acquisition of time-resolved fluorescence signals for only one cell did not give a sufficient signal/noise ratio for further analysis, the signals were summed over measurements repeated for five independent cells induced to either state1 or state2. Figure S4 shows the results for 10 cells, in which five cells were induced to state1 (upper panels), and the others induced to state2 (lower panels). Figure 3 shows the time-resolved fluorescence two-dimensional (2-D) maps in the PSII-rich (AD) and PSI-rich (BE) domains averaged over five cells induced to state1 (AB) and state2 (DE). Figure 3CF shows the time-integrated spectra whose peak heights at around 680 nm were normalized to unity. The spectra of the PSI-rich domains (red lines) show higher relative intensities of the PSI bands at around 710 nm for both state1 and state2 cells, as expected. These spectral features are typical to ST-operating *Chlamydomonas* cells, confirming the effective induction of ST with the present method.

**Figure 3.**
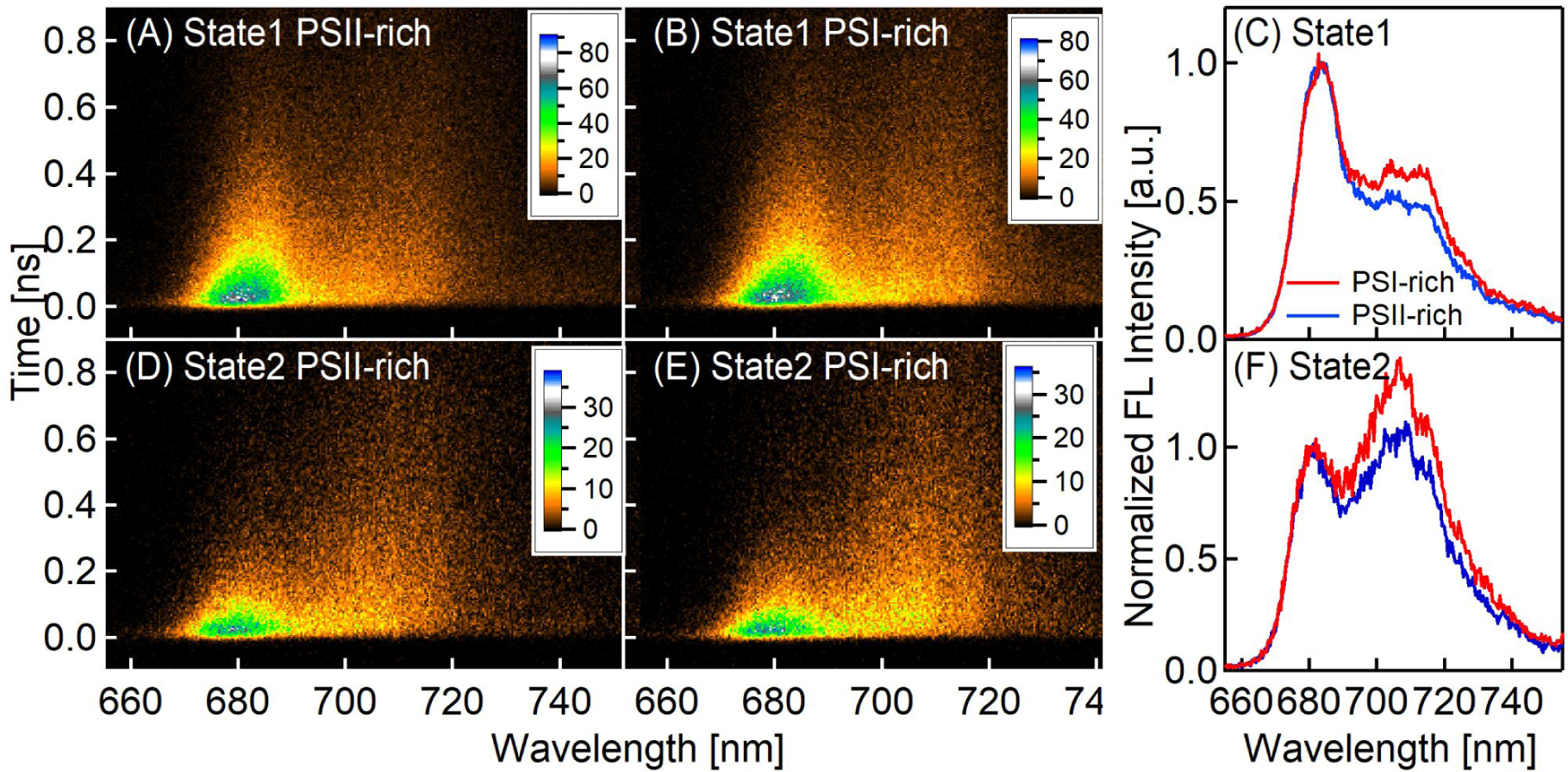
Streak images for the specific microscopic areas of Chlamydomonas cells measured with the cryogenic streak-camera optical microscope system. Each obtained streak image is accumulated for five measured cells induced to state1 ((A) for PSII-rich and (B) for PSI-rich regions) or state2 ((D) for PSII-rich and (E) for PSI-rich regions). Blue and red solid lines in panels (C) (for state1) and (F) (for state2) represent steady state fluorescence spectra for PSII-rich and PSI-rich regions, respectively, calculated by integrating the streak images (A, B, D, E) along the time axis.

To derive insight into the EET dynamics, we conducted the global fitting of the time-resolved fluorescence spectra. The 2-D maps in Fig. 3ABDE were divided into 17 segments along the horizontal wavelength axis. Integrating the signal in each segment along the wavelength axis gives the fluorescence time profile in the given wavelength region (the blue lines in Fig. S5). The curves were fitted to the sum of four exponential decay curves (the red lines in Fig. S5) convolved with the IRF. In the present study, the IRF was approximated by a Gauss function with a full width at half maximum (FWHM) of 24 ps. This value was obtained by averaging the FWHM values of the Gaussians that fit the MG time profiles at five different wavelengths (see Fig. S3A). The fluorescence time profiles were fitted globally with the constraint that each time constant has a common value between all of the fluorescence kinetics at different wavelengths. The amplitude of each exponential component plotted against the wavelength is called the fluorescence decay-associated spectra (FDAS). The positive and negative amplitudes of an exponential fluorescence kinetics correspond to the decaying (energy-donating) and rising (energy-accepting) kinetics, respectively.(Lofroth, 1986) Therefore, a pattern with positive and negative bands at shorter and longer wavelength sides of the FDAS, respectively, is an indication of a downhill EET with the time constant of that component.

Figure 4 shows the thus-obtained FDAS of the PSII-rich (left panels) and PSI-rich (right panels) domains in the state1 cells (upper panels) and the state2 cells (lower panels). The fastest FDAS components in all panels in Fig. 4 exhibit signs of the EET, characterized by the positive and negative bands at around 670 nm and 680–685 nm, respectively. These components probably reflect the EET from the peripheral antenna mainly to PS core complexes in state1 and state2 cells, respectively. This interpretation is consistent with the fact that the present excitation wavelength at 460 nm is preferentially absorbed by carotenoids and Chl-*b*s, the latter of which are bound only to peripheral antennas. The EET from Chl-*b* to Chl-*a* within the peripheral antenna might be too fast to be resolved by the present setup. For state2 cells, the second-fastest FDAS components also showed EET features represented by a broad negative band at around 710 nm. This may reflect the EET to the red Chls in PSI. A recent study of PSI core and PSI-LHCI supercomplexes isolated from *Chlamydomonas* suggested that the red Chls in *Chlamydomonas* are bound to LHCI but probably not to the PSI core complex.(Giera et al., 2018) Thus, the negative band of the second-fastest FDAS of state2 cells reflects the EET (more precisely the energy-equilibration) kinetics to the red Chls in LHCI. The third and fourth FDAS components are positive over the whole spectral range. The former and latter components mainly reflect the fluorescence decay kinetics of PSII and PSI, respectively; nonetheless, they also have mixed contributions from both PSs.

**Figure 4.**
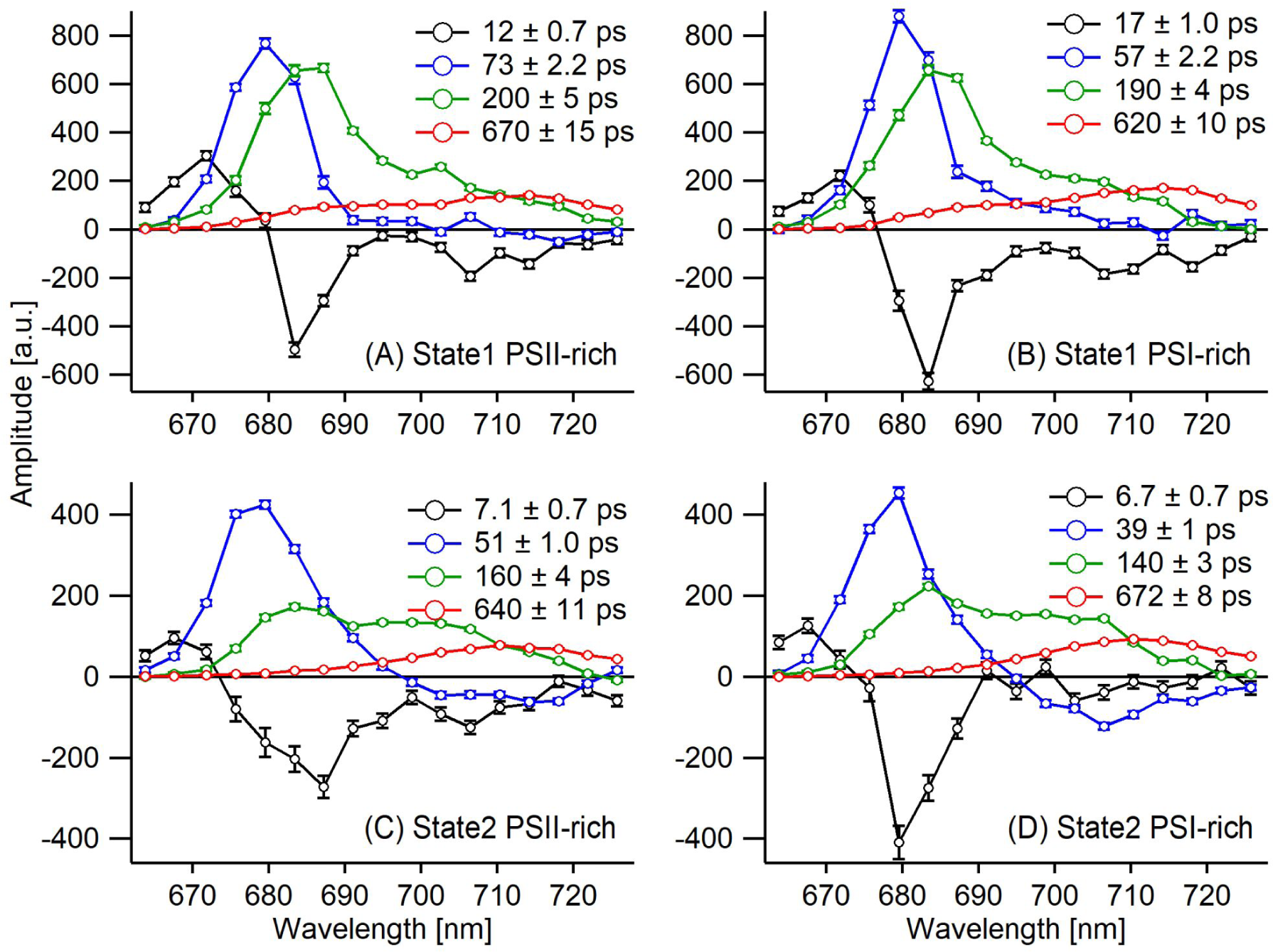
Decay-associated spectra calculated for the intracellular PSII-rich (left) and PSI-rich (right) regions in state1 (top, A and B) and state2 (bottom, C and D). Black, sky-blue, green, and red curves show the spectra associated with the indicated time constants. The error bars and values to the right of ± symbols in the annotation indicate the standard error of the global fitting with the exponential decay model.

Finally, we show data in Fig. S6 that ensures the ability of the present setup to resolve the fluorescence decay dynamics with time constants less than the FWHM of the instrumental response time (24 ps). To this end, we measured MG solutions in water-glycerol mixtures with two different glycerol contents. It has been reported that the fluorescence decay time of the emission from its S_1_ state slows with the enhanced viscosity of the solvent.(Yoshizawa et al., 1998) Yoshizawa et al. obtained a simple relation between the time constant *τ* and the solvent viscosity *η*, expressed as *τ* ∝ *ƞ*^2⁄3^. We prepared MG solutions in water-glycerol mixtures with glycerol volume ratios of 65.3% and 76.7%. The viscosities of the former and latter mixtures were calculated to be 19.5 cP and 54.5 cP, respectively, based on a reported formula.(Cheng, 2008) The above *τ* ∝ *ƞ*^2⁄3^ relation predicts that the 76.7% glycerol volume ratio gives an MG S_1_ decay time exactly two-times slower than the 65.3% glycerol volume ratio. Figure S6 shows the fluorescence decay kinetics of the MG S_1_ emissions in these two mixed solvents together with that of the MG aqueous solution as the instrumental response function. The fitting of the decay kinetics gave time constants of 9 ps and 20 ps for the 65.3% and 76.7% glycerol volume ratio samples, respectively. Two independent measurements reproduced the results. The obtained time constants demonstrated an excellent agreement with the relation reported by Yoshizawa et al. Thus, we could safely confirm the ability of our setup to resolve the fluorescence kinetics with a time constant of ca. 10-ps.

## Discussion

In our previous studies, we measured the intracellular local fluorescence spectra of photosynthetic organisms at 80 K.(Fujita et al., 2018; Fujita et al., 2022) Although this has been a powerful technique for identifying intracellular PSII-rich and PSI-rich domains, there has been ambiguity in the interpretation of the results. Since we could detect the fluorescence decay at only one single wavelength with the setup used in the previous study, we could not determine whether the rapid decay of fluorescence was due to the EET from the peripheral antenna to the core complex or the efficient quenching of the protective function. We could spatially resolve the PSII-rich and PSI-rich domains based on the intensity of the fluorescence band specific to each PS. However, we could not determine whether the relatively high local PSII fluorescence comes from the higher local PSII concentration (Fig. 1A) or the locally enlarged antenna size of PSII (Fig. 1B). The former interpretation is consistent with several reports based on electron microscopic observations that indicated a certain level of segregation of PSII and PSI.(Vallon et al., 1985; Wietrzynski et al., 2020) However, it is still difficult to reject the latter interpretation attributing the inhomogeneous PSII/PSI fluorescence ratios to the intracellular inhomogeneity of the antenna sizes of PSs.

The present study provides a key clue as to resolving the ambiguity cited above. The global analysis, providing the FDAS, gave information on the EET dynamics in each of the PSI-rich and PSII-rich domains. If the higher local PSII/PSI fluorescence ratio comes from the larger PSII antenna size, the EET from the peripheral antenna to the PSII core complex should take more time due to the elongated EET pathway. Thus, we will be able to distinguish the two situations depicted in Fig. 1. However, we must point out here the remaining limitation of the present measurement. Although the intracellular PSI-rich and PSII-rich domains could be spatially resolved with our setup, the fluorescence dynamics cannot be interpreted as pure PSI and PSII signals. They still contain mixed contributions from PSI and PSII with a different relative weight of each component. Therefore, we should interpret each FDAS component with careful consideration of the above-mentioned limitation.

The EET process is reflected in an FDAS with a positive band accompanied by a negative dip on its longer wavelength side. In Fig. 4, we can find one FDAS components with the EET-specific characteristic in state1 cells, whereas there are two in state2 cells: the fastest FDAS in both the state1 and state2 cells, and the second fastest FDAS in the state2 cells. In state1, the majority of LHCII binds to PSII. Therefore, it is reasonable to interpret that the fastest FDASs of state1 cells reflect mainly the EET from LHC to PSII in both PSI-rich and PSII-rich domains. This interpretation is consistent with the time-integrated spectra of state1 cells shown in Fig. 3(C). They have much stronger contributions of PSII than PSI for both the PSI-rich and PSII-rich domains. Several previous reports(Snellenburg et al., 2017; Giera et al., 2018) have suggested that the EET dynamics in the PSI-LHCI super-complex is much faster (8 ps) than the time constants of the fastest FDAS of the state1 cells. Therefore, we think that the EET in PSI does not significantly contribute to the fastest FDAS of state1 cells. Importantly, the time constant of the fastest FDAS of state1 cells for the PSI-rich domain is even longer than for the PSII-rich domain. This clearly demonstrates that the higher PSII fluorescence bands of the PSII-rich domains in state1 cells are not due to the larger antenna size of PSII but are due to the larger PSII concentration instead.

In state2, we found two EET-type FDASs, the fastest and second fastest ones. The profiles of the fastest FDASs in state2 cells are similar to those in state1 cells. Accordingly, we assign this component mainly to the EET from the LHC to PSII core complex. Their time constants significantly decreased, from 12-17 ps in state1 to ca. 7 ps in state2. This shortening of the EET time constant is consistent with the above-mentioned assignment because it can be interpreted as reflecting the detachment of the peripheral antenna from PSII upon the transition to state2. The time constant of the fastest FDAS in the state2 cells showed negligible differences between the PSII-rich and PSI-rich domains, demonstrating that this light-harvesting dynamics are not significantly different between the PSII-rich and PSI-rich domains. This again suggests that the antenna sizes of the PSII in the PSI-rich and PSII-rich domains are similar to each other in state2. The negative dips at around 710 nm of the second fastest FDAS are assigned to the rise of emission from the red Chls, which have been reported to be located only in LHCI but not in the PSI core in *Chlamydomonas*.(Giera et al., 2018) These negative dips became visible only after the transition to state2. Accordingly, we assign the second fastest FDAS of the state2 cells to the excitation-energy equilibration in PSI-LHCI-LHCII super-complexes that emerged after the state1 to state2 conversion. The time constant of the second fastest FDAS component for the PSI-rich domain was even faster than that for the PSII-rich domain. This also denies that the antenna size of PSI is larger in PSI-rich domains than in PSII-rich domains. The negative dip at 710 nm was deeper for the PSI-rich than for the PSII-rich domain. This can be regarded as an indication of the enhanced PSI concentration in the PSI-rich domain.

In comparisons of the EET-type FDAS components between the PSI-rich and PSII-rich domains, we detected no evidence supporting the situation depicted in Fig. 1(B). Thus, we conclude here that the inhomogeneous antenna sizes are not the main origin of the inhomogeneous PSII/PSI fluorescence ratio. The present study supports the interpretation adopted in our previous studies,(Fujita et al., 2018; Fujita et al., 2022) that the inhomogeneous PSII/PSI fluorescence ratios within single cells reflect the concentration inhomogeneity of the PSs.

Here, we discuss the time constant of the second-fastest FDAS in state2 cells. They are assigned to the excitation-energy equilibration in the red Chl-containing super-complex (PSI-LHCI-LHCII) as described above. It is conspicuous that the time constants of 40 to 50 ps for these second-fastest FDASs in state2 cells are much slower than those reported so far for the isolated PSI-LHCI-LHCII super-complex. The EET-type FDAS was reported for the isolated PSI-LHCI-LHCII super-complex of *Chlamydomonas*.(Le Quiniou et al., 2015) Its electron micrograph clearly indicated the binding of two additional LHCII trimers. Although its fluorescence dynamics was measured at room temperature, the EET-type FDAS reported for this isolated super-complex has a time constant of 1.8-3.5 ps, which was one order of magnitude faster than that reported here. On the other hand, Wlodarczyk et al. have reported slower EET, or energy-equilibrium, dynamics to the red Chls of LHCI in vivo.(Wlodarczyk et al., 2016) They used a PSII-deficient *Chlamydomonas* strain to analyze the PSI-LHCI-LHCII super-complex in vivo. They reported two time-constants, 26 and 113 ps, of the EET-type FDAS. Although these were not exactly the same as, but more or less consistent with, those obtained in our present study. Taken together, the EET to the red Chls in LHCI takes much more time in intact state2 cells than in the isolated PSI-LHCI-LHCII. This indicates that the antenna size of the PSI super-complexes in the intact cells induced to state2 is drastically more enlarged than in the isolated PSI-LHCI-LHCII super-complex.

The present study revealed distinct properties of FDASs between the PSII-rich and PSI-rich domains. However, the difference was not so evident, especially for state1 cells. We notice here a difference in the spectral profiles of the fastest FDAS components between PSII- and PSI-rich domains of state2 cells. The fastest FDAS in the PSI-rich domain of state2 cells shows a negative amplitude that is significantly reduced over the wavelength range from 690 to 720 nm. This is a conspicuous feature of this FDAS component as compared with the fastest FDAS shown in the other three panels (ABC) of Fig. 4, which were negative over a wide range from 675 to 720 nm. We assigned the fastest FDAS mainly to the EET from the peripheral LHC to PSII core. Accordingly, the negative band in the 690 to 720 nm wavelength region can be assigned to the temporal rise of the vibronic fluorescence band of the PSII core. It is unusual that the fastest FDAS has a negligible negative sign in the 690-720 nm region only for the PSI-rich domains of state2 cells. One possible explanation for the absence of negative amplitude in the fastest 6.7-ps FDAS is that the rising kinetics are canceled out due to the overlap of the decaying kinetics with a similar time scale.

Here, we refer to our previous observation on the same *Chlamydomonas* samples by the cryo-FLIM technique,(Fujita et al., 2022) which might be relevant to the absence of the negative band in the fastest FDAS, as described above. In that study, we detected fluorescence decay kinetics only at around 680 nm, assigned mainly to LHCII and partially to the PSII core complex. The excitation wavelength was at 460 nm, the same as that of the present study. In that study, we found that PSI-rich domains in state2 cells have some specific parts that accumulate a characteristic component with an extremely rapid fluorescence decay at 680 nm with a time constant of less than ca. 3 ps. An unusual feature in the rapid fluorescence dynamics in the PSI-rich domains of the state2 cell was again detected in the present study as described above. We interpreted the above-mentioned observation in the previous study as a suggestion of the accumulation of highly quenched LHCII aggregates in the PSI-rich domains in state2 cells. This interpretation relied on the assumption that a part of LHCII detached from PSII upon conversion to state2 may fail to bind to PSI and form the quenched state. Here, we tentatively interpret the absence of the negative sign in the 690-720 nm region of the fastest FDAS in the PSI-rich domains of the state2 cells as follows: the fluorescence rise in the 690-720 nm spectral region was cancelled by the overlap of the rapid decay of the quenched LHCII with a red-shifted emission band. To confirm this speculative explanation, we should further enhance the S/N ratio of the data.

## Conclusions

We developed a cryo-microscopic system connected to a streak camera. This enabled us to measure the time-resolved fluorescence spectra in either the intracellular PSII-rich or PSI-rich domains within single cells. The rate constants of the EET from LHC to PSII and to PSI deduced from the FDAS analysis are essentially the same between the apparent PSII-rich and PSI-rich regions. Based on this result, we suggested that the higher/lower PSII/PSI fluorescence ratio comes from the enlarged/reduced local PSII concentration, not from the larger/smaller antenna size of PSII. Thus, the intracellular inhomogeneity in the PSII/PSI fluorescence intensity reflects the inhomogeneous relative concentrations of the PSs. Further, it is worth noting that the present study also suggests that PSIs in intact cells have larger antenna sizes than those of isolated PSI containing super-complexes. This was concluded from the slower time constant of the EET to the red Chls in LHCI estimated for the *Chlamydomonas* cells than for the previously reported isolated PSI-LHCI-LHCII.(Wlodarczyk et al., 2016)

This technical advancement reported in the present study may provide a powerful new tool for analyzing various research targets. Since the light-harvesting dynamics in natural photosynthetic systems are generally dependent on the position in a cell and on the environmental conditions surrounding the cell,(Pinnola et al., 2015; Ruban, 2016) they will be suitable targets of the methodology developed. The technique will be also applied efficiently to the study of the reaction dynamics of isolated chromoproteins at the single-molecule level.(Kondo et al., 2017; Squires and Moerner, 2017; Brotosudarmo et al., 2022) Various artificial photosynthetic systems will be the potential research targets. For example, the method will be effective in revealing spatial inhomogeneity in the EET dynamics of various pigment-aggregate-based artificial light-harvesting systems,(Bösch et al., 2016; Shoji et al., 2016; Matsubara and Tamiaki, 2019) nanoparticle-biomaterial hybrid-based systems.(Noji et al., 2011; Kawahara et al., 2020)

## Materials and Methods

### Sample preparation

*Chlamydomonas reinhardtii* cells (strain 137c) were cultured as described previously.(Fujita et al., 2018; Zhang et al., 2021; Fujita et al., 2022; Zhang et al., 2022) Briefly, cells cultured on a TAP (tris-acetate-phosphate-containing) agar plate were transferred to a TAP aqueous medium and cultured for three or four days under white light from an LED (50 μmol photons m^−2^s^−1^). Then, the cells were transferred to a photoautotrophic condition in a high-salt (HS) aqueous medium one or two days before the experiment. The cell suspension in the HS medium was packed in a home-made copper sample holder with two quartz windows that were 0.3 mm thick. The sample holder was connected to the cold head of the cryostat (Microstat, Oxford Instruments, Eynsham) and cooled to 80 K. For the induction of cells to state1, the cell suspension set in the microscope was irradiated with light at 710 nm (PSI light, 150 μmol photons m^−2^s^−1^) for 15 min at 23°C. State2 induction was carried out by incubating the cells under an anoxygenic environment: the cell suspension was supplied with glucose (20 mM) and glucose oxidase (50 U mL^−1^) and then immediately packed in the sample holder and incubated in the dark for 15 min. After induction to state1 or state2, a liquid nitrogen flow was started to initiate cooling.

### The setup of cryogenic optical microscopy connected with a streak camera

We extended our cryogenic microscopic system to detect time-resolved fluorescence spectra of the local PSI-rich and PSII-rich domains within single *Chlamydomonas* cells. The excitation beam was provided from a sub-picosecond pulsed (80 MHz) Ti:S (MAITAI, Spectra-Physics, Mountain View, CA) converted to the second harmonics at 460 nm by a BBO (β-BaB_2_O_4_) crystal. The typical excitation power was ca. 100 nW at the sample position. A beam-splitter cube separates the fluorescence light (Fig. S1) into two paths: the 30% transmitted through the cube was input to the polychromator equipped with the CCD camera for ordinary spectral detection, and the reflected 70% was focused to the optical-fiber bundle whose exit was coupled with the optical system of the streak camera. The confocal pinhole was set before the beam-splitter cube, ensuring the observations of the same focal plane on the sample for the CCD and streak-camera channels.

For an actual measurement, we first found a cell by a transmission image, and then we obtained the spectral image of the target cell by the raster scanning of the laser with the Galvanic mirror pair (Figs. S1, S2a). All of the pixels in the image obtained by this preliminary scanning contain fluorescence spectra of the corresponding local points in the cell. For each pixel, the following two parameters were calculated:

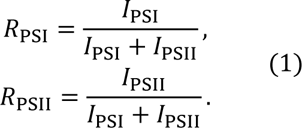

Here, *I*_PSI_ and *I*_PSII_ are the integrated spectral intensity over the PSI (706-727 nm) and PSII (676-692 nm) spectral ranges, respectively. Then, PSI-rich and PSII-rich pixels were selected as those in which *R*_PSI_ and *R*_PSII_ exceed certain pre-defined threshold values, respectively. In order to measure the time-resolved fluorescence spectra of thus-extracted PSI/PSII-rich domains, the laser spot was scanned only over the PSI/PSII-rich domains. The sampling of the streak camera was carried out during domain-selective scanning. The above sequence of measurements was conducted by our self-written control program (LabView, National Instruments, Austin, TX) of the microscopic system (Fig. S2).

Since the fluorescence light detected by the streak camera passed through an optical fiber bundle, its arrival time was delayed more at a shorter wavelength, due to the higher refractive index of the fiber material. To correct this dispersion effect, we measured the fluorescence time profile of an aqueous solution of a dye, malachite green (MG), with a concentration of 10^−4^ M with the current setup at room temperature. An aqueous solution of MG is known to show very rapid fluorescence decay (less than 1-ps lifetime) with a broad spectral width at room temperature.(Yoshizawa et al., 1998) It can be a suitable standard sample for estimating both the instrumental response function (IRF) and the wavelength-dependent delay effect. Figure S3 shows the time profile of an MG aqueous solution measured by the current setup. The peak position along the time axis was obviously delayed at shorter wavelengths. The time profiles at several wavelengths (indicated by crosses in panel B) were fitted to a Gauss function (panel A). Then, the wavelength dependence of the peak position along the time axis was fitted to a straight line (red line in panel B). The slope of the linear function was estimated to be −1.0033 ps/nm and was used to correct the dispersion effect. Figure S3C shows the corrected time profile of the MG solution, showing a horizontal shape—at least in the spectral region of interest (660–740 nm)—as expected.

Supporting Information. Additional figures of experimental set-up and data as discussed in the text.

## Acknowledgements

We thank Professor Jun Minagawa and Dr. Ryutaro Tokutsu at the National Institute for Basic Biology for the kind gift of PSII-LHCII and PSI-LHCI samples isolated from *Chlamydomonas*.

## Author Contributions

Y.S. conceived the experiments; Y.F. performed the experiments and analyzed the data. Y.F. and Y.S. wrote the manuscript with contributions of all authors; X.Z and S.Y. supervised the experiments and writing process. Y.S. agrees to serve as the author responsible for contact and ensures communication.

